# Towards Generalizability and Robustness in Biological Object Detection in Electron Microscopy Images

**DOI:** 10.1101/2023.11.27.568889

**Authors:** Katya Giannios, Abhishek Chaurasia, Cecilia Bueno, Jessica L. Riesterer, Lucas Pagano, Terence P. Lo, Guillaume Thibault, Joe W. Gray, Xubo Song, Bambi DeLaRosa

**Affiliations:** Micron Technology, Inc, Boise, 83707, ID, USA.; Oregon Health & Science University, Portland, 97239, OR, USA.; Chaos Industries, Bellevue, 98004, WA, USA.

**Keywords:** Group normalization, data augmentation, robustness, generalization, electron microscopy, object detection

## Abstract

Machine learning approaches have the potential for meaningful impact in the biomedical field. However, there are often challenges unique to biomedical data that prohibits the adoption of these innovations. For example, limited data, data volatility, and data shifts all compromise model robustness and generalizability. Without proper tuning and data management, deploying machine learning models in the presence of unaccounted for corruptions leads to reduced or misleading performance. This study explores techniques to enhance model generalizability through iterative adjustments. Specifically, we investigate a detection tasks using electron microscopy images and compare models trained with different normalization and augmentation techniques. We found that models trained with Group Normalization or texture data augmentation outperform other normalization techniques and classical data augmentation, enabling them to learn more generalized features. These improvements persist even when models are trained and tested on disjoint datasets acquired through diverse data acquisition protocols. Results hold true for transformer- and convolution-based detection architectures. The experiments show an impressive 29% boost in average precision, indicating significant enhancements in the model’s generalizibality. This underscores the models’ capacity to effectively adapt to diverse datasets and demonstrates their increased resilience in real-world applications.

---

In the biomedical field, deep learning approaches have shown promising results in tasks such as image classification, segmentation, and detection. However, their practical utility is limited by their poor generalization capability. To achieve broad adoption of machine learning (ML) in biomedical applications, robust and generalizable models are essential.

One of the main challenges is the lack of diverse and representative datasets. Many publicly available biomedical datasets are limited to specific experiments, tissue types, or acquisition instruments. Even a large amount of data may lack diversity in terms of subject population or data collection protocols, creating significant gaps in the data distribution. Consequently, when applying state-of-the-art machine learning models trained on these datasets to external data, significant performance degradation is observed [1, 2]. In addition to dataset limitations, data shifts pose another challenge. Changes in data collection protocols or variations in behavior of specimens can lead to shifts in the data distribution [3]. Moreover, after deploying a model, the observed data distribution may change over time, deviating from the original training data distribution. These unavoidable shifts can result in a significant drop in model performance, often exceeding 10% [4]. We aim to address this disparity and propose improvements to mitigate the impact of data distribution shifts.

There are several ways to tackle the challenges posed by the lack of diverse training data. One of the most common approaches is to collect more data. However, in the biomedical field this is often not feasible due to limitations in the availability of specimens. Another possibility is to use transfer learning. Transfer learning involves pre-training a model on a large and diverse dataset, like ImageNet [5], and then fine-tuning it for the specific biomedical task. By leveraging knowledge from the pre-training, the model can adapt to the new problem effectively. For effective transfer learning, it is essential to have a substantial number of samples, a wide range of diverse images, and a close resemblance between the training data and the target application [6]. Yet another approach towards generalizability is Domain Adaptation. Domain adaptation focuses on minimizing the differences between the training and test domains, allowing models to generalize better. Various techniques have been developed to align or adapt the learned representations to the target domain. An extensive survey of different categories of domain adaptation methods for medical images is given by Guan in [7]. Prior knowledge of the domain differences is typically required, and these techniques often involve complex modeling and adaptation processes.

Hendrycks and Dietterich [8] highlighted the limited generalizability of classification models trained on ImageNet when tested on a distorted variant called ImageNet-C. Michaelis, Mitzkus, and Geirhos [9] demonstrated similar findings on the task of object detection. Additional benchmarking studies conducted by [9–11] have further emphasized this issue. These studies collectively suggest that augmenting the training data with various levels of image corruption can enhance the robustness of models and improve their generalization capabilities. Xu and Mannor [12] emphasized the importance of robustness as a key characteristic for learning algorithms to achieve effective generalization. While data augmentation has the potential to improve generalization performance, training models solely on handpicked corruptions can lead to memorization of those specific corruptions, limiting the model’s ability to generalize to new ones [13].

In this study, the objective is to propose a recipe that utilizes simple existing techniques to enhance the robustness and generalizability of biomedical data. We explore the factors contributing to performance degradation, with a focus on texture analysis to capture dataset variability and its impact on machine learning algorithms. Extensive experiments are conducted using electron microscopy (EM) images, leading to the following conclusions: i.) generalization improves significantly when texture-specific information is removed by transitioning from batch normalization to image-specific normalization processes, and ii.) fine-tuning combined with data augmentation techniques specifically targeting the enrichment of image textures enhances overall model robustness.

We demonstrate our findings with a detection task using EM images. Accurate identification of organelles is crucial for analyzing cellular structures, and our specific focus is on the automated detection of mitochondria. The dataset utilized for this task comprises EM images accompanied by annotations in the form of manually-drawn bounding boxes around each instance of the organelle (refer to Fig. 1**a**). We examined four datasets, including one private dataset and three publicly available ones:

**Fig. 1.**
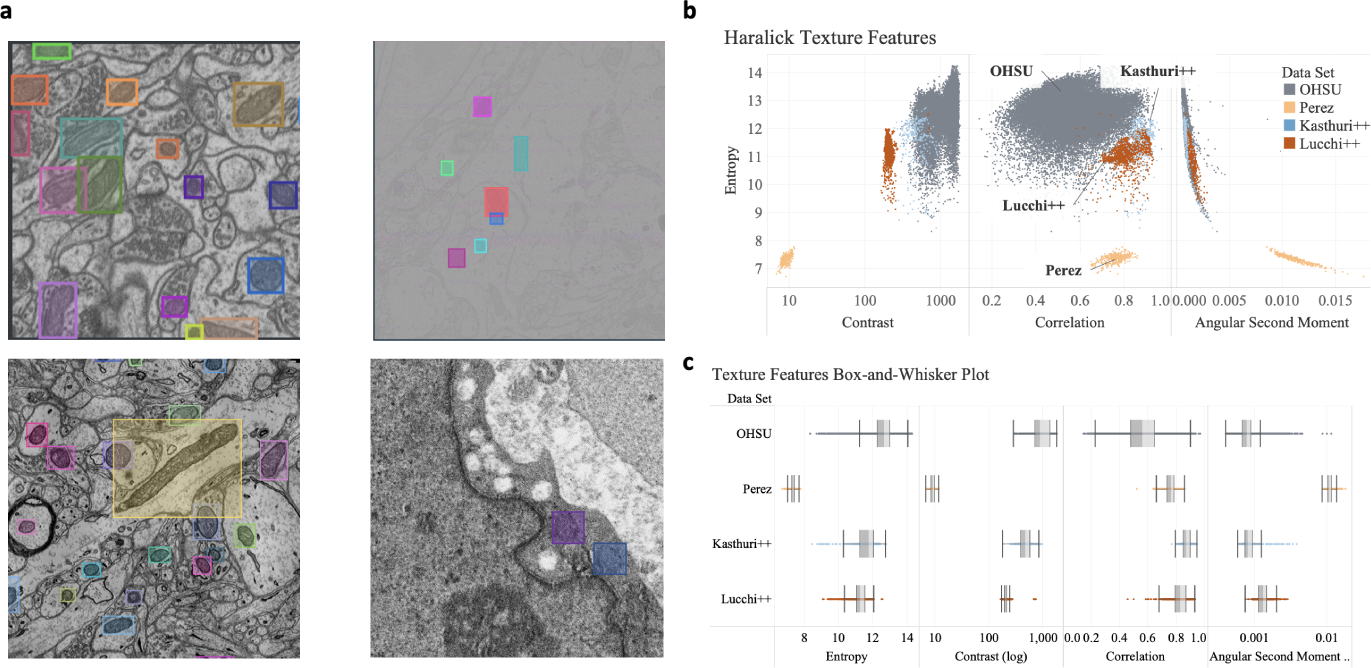
Towards generalizing and robust detectors: **a.** Selected images from (clockwise) Lucchi++, Perez, OHSU, Kasthuri++ datasets; **b.** Haralick texture spatial distributions of the datasets highlighting textural variability observed in the mitochondria; **c.** Box-whisker plot for four of the Haralick features for the different datasets.

- *Private dataset: Oregon Health & Science University (OHSU) Dataset.* The OHSU dataset comprises 17 samples obtained from stage-three and -four metastatic cancer patients, covering various types of cancers such as breast, pancreas, ovarian, and liver originating from different body locations. The data collection process adhered to a rigorous protocol outlined in [14]. Scanning electron microscopy (SEM) was employed for data acquisition, with large-format montages utilized. Each individual tile within the montages measured 6000 x 4000 pixels in size, with a resolution of 4nm per pixel. During tile acquisition, a 10% overlap was maintained. The images in the dataset have been annotated for the detection of mitochondria, endosomes, nuclei, and nucleoli.
- *Public dataset: Kasthuri++, Lucchi++, Perez.* The three public datasets consist solely of EM images obtained from mouse brain tissue. These images were captured using a range of SEM-based technologies. Initially, these datasets were published for the purpose of 2D or 3D segmentation tasks. For our study, we transformed the segmentation masks into bounding boxes specifically for detection purposes, and they were exclusively employed in 2D detection inference.
  – Kasthuri++ [15]: The images were acquired using Automated Tape-Collecting Ultramicrotome ribbons of serial sections and SEM (ATUM-SEM). The tissue samples were obtained from dense mammalian neuropil tissue, specifically from layers 4 and 5 of the S1 primary somatosensory cortex. The original dataset was divided into two stacks: one for training and the other for testing. For our experiments, we exclusively utilized the single images from the test stack. The dimensions of the test stack are 1334 *×* 1553 *×* 75 voxels, with a voxel size of 3 *×* 3 *×* 30 nm. The initial release of the data was made publicly available by [16], and subsequent re-annotation was performed by [15].
  – Lucchi++ [15]: The serial images in the Lucchi++ dataset were obtained using focused ion beam-scanning electron microscopy (FIB-SEM). The images were acquired from a 5 *×* 5 *×* 5*µm* section of the hippocampus of a mouse brain, with a voxel size of 5 *×* 5 *×* 5 nm. The dataset was divided into training and testing sets, each comprising the first 165 slices of a 1065 *×* 2048 *×* 1536 image stack. The dataset includes manually created mitochondria segmentation masks. Initially, it was published as the EPFL Hippocampus Data [17], and later the image annotations were revised, corrected, and republished as the Lucchi++ dataset. For our experiments, we only utilize the 165 2D test images from this dataset.
  – Perez [18]: In the Perez dataset, the images were obtained using serial block face-scanning electron microscopy from a 124.8 *×* 93.6 *×* 38.5*µm* section of the hypothalamus of a mouse brain. The voxel size for these images is 7 *×* 7 *×* 30 nm. It is important to note that the test images in the Perez dataset do not have labels for all instances of mitochondria, leading to a significantly lower precision in the detection task.

We selected six prominent detection architectures from the Open MMLab Detection Toolbox repository [19]: Deformable DETR (DDETR) [20], DETR [21], Dynamic R-CNN [22], TOOD [23], Sparse R-CNN [24], and YOLOv3 [25]. Among the selected architectures, Dynamic R-CNN, TOOD, and Sparse R-CNN belong to the family of Region-Based Convolutional Neural Networks (R-CNNs). These R-CNNs typically operate in two stages and are commonly used for instance segmentation. On the other hand, YOLOv3 and the two transformer-based architectures (DETR and DDETR) perform instance segmentation in a single stage. DETR and DDETR are transformer-based architectures, while the remaining networks are convolutional neural networks. More comprehensive details regarding the architectures and reproducibility can be found in the Methods section.

Each of the selected networks boasted state-of-the-art performance on specific datasets at the time of their publication. Furthermore, each network has its own distinct advantages in various aspects, such as the availability of computational resources for training, the ability to detect small objects effectively, or fast inference speed. In terms of performance evaluation, we utilize the Average Precision (AP) metric, which is commonly used in the MS COCO challenge [26]. Specifically, we focus on the average precision for bounding box predictions with an Intersection over Union (IoU) of at least 50% (AP50). Detailed information about these performance measures can be found in the Methods section.

In our study, we divided the OHSU dataset into training, validation, and test datasets and trained the six selected networks on this data. Subsequently, we evaluated the performance of these trained networks on the test datasets of the public datasets (Kasthuri++, Lucchi++, Perez), as well as on the OHSU test dataset. As expected, the results demonstrated significant variation in average precision across the different methods. Moreover, we observed variations in the transferability of these methods to unseen datasets, as presented in Table 1.

**Table 1.** Establishing Baseline: Average Precision 50 (AP50) of the six models (all trained on OHSU data) when generalizing to four different test sets.

| Methods | OHSU DS | Kasthuri++ | Lucchi++ | Perez |
| --- | --- | --- | --- | --- |
| DDETR | 56.6% | 51.4% | 56.7% | 16.3% |
| DETR | 60.0% | 51.5% | 58.7% | 0.9% |
| Sparse RCNN | 54.9% | 37.0% | 35.9% | 0.0% |
| TOOD | 54.7% | 4.7% | 48.3% | 0.2% |
| Dynamic RCNN | 49.5% | 17.1% | 43.3% | 0.0% |
| YOLOv3 | 53.8% | 36.2% | 44.0% | 4.8% |

The two transformer-based methods, DDETR and DETR, exhibited relatively good generalization on the Lucchi++ dataset, with performance levels close to the AP50 achieved on the OHSU test dataset (which shares the same data source as the training data). Additionally, these methods achieved reasonable average precision on the Kasthuri++ dataset. However, their performance was notably poor on the Perez dataset.

The performance of the three R-CNN-based networks showed significant variability on the Lucchi++ dataset, ranging from 35.9% to 48.3% in terms of average precision. Their performance further declined on the Kasthuri++ dataset and was nearly negligible on the Perez dataset, likely due to the low contrast in the images. Similar trends were observed for the YOLO network, with comparable performance patterns across the different datasets.

Upon further analysis of the data and its texture characteristics, we have observed a significant difference in the contrast-entropy distribution among the datasets, as depicted in Figure 1 **b-c**. This distinction is not only evident when comparing the public datasets to the training OHSU data but also within the OHSU dataset itself. This observation presents an opportunity to enhance the overall performance and generalizability of the models. The primary objective of this paper is to explore available techniques, which are not commonly adopted, for improving the domain generalizability of the models. From the results presented in Table 1, it is evident that none of the models demonstrate satisfactory generalization across all datasets. Despite the presence of semantically similar structures and the shared nature of EM images, the variability introduced by different tissue types, collection protocols, and instruments creates challenges for detection across datasets. Here, we push forward the notion that the variability lies within the texture of the images, and we aim to investigate and find approaches to “normalize” this texture variability, ultimately leading to improved overall performance.

Our contributions include:

- A comprehensive investigation into the factors contributing to the performance drop, focusing on the data variability captured through texture analysis.
- Quantification of the impact of deepening the backbone architecture, replacing it with a more robust backbone, or utilizing pre-training techniques such as stylized ImageNet or unsupervised pre-training on biomedical-relevant data.
- Introducing texture augmentation as a means to incorporate additional plausible texture variability in the training data.
- Evaluation of the effect of replacing batch normalization in the backbone with group normalization, thereby mitigating the influence of contrast variability.

These contributions aim to shed light on the underlying causes of performance limitations and provide potential solutions to improve the robustness and generalizability of the models.

## 1 Results

In our analysis, we employ two tools: Haralick’s texture features and tdistributed stochastic neighbor embedding (t-SNE) [27].

Haralick’s texture features [28] comprise a set of textural features calculated from the co-occurrence matrix representation of the original image. They provide valuable insights into the texture characteristics of the images, allowing us to quantify and compare textural variations across different datasets.

To further explore the internal representation of the images by the detectors, we utilize t-SNE, which is a dimensionality reduction technique. T-SNE enables us to visualize the high-dimensional representations of the images in a lower-dimensional space, providing a better understanding of the relationships and similarities between images as perceived by the detectors.

Texture in image processing refers to the spatial variation of brightness intensity among pixels. To quantify and evaluate the texture differences within and between datasets, we utilize Haralick’s texture features [28]. These features are computed from a Gray Level Co-occurrence Matrix (GLCM), which captures the co-occurrence of neighboring gray levels in the image for a specified offset. The GLCM is a square matrix with dimensions equal to the number of gray levels (typically 255) in the region of interest. It records the normalized frequencies or probabilities of each combination of gray level co-occurrences in the image. From the GLCM, we can calculate 13 Haralick features that characterize the texture properties of the image. As a consequence of a high level of correlation among the 13 features we follow the approach of [29] and focus on four specific Haralick features: sum entropy, correlation, contrast, and angular second moment. The entropy feature measures the level of randomness or information content in the intensity distribution of the image. A higher entropy value indicates a more diverse and complex texture. On the other hand, correlation measures the linearity or presence of linear structures in the image. Contrast quantifies the local variations in intensity within the image. A higher contrast value indicates pronounced difference between neighboring pixel intensities.The angular second moment reflects the uniformity of the image texture. Notably that EM images inherently contain noise, which can influence the magnitude of the Haralick features [30]. While denoising the images before the analysis could be beneficial, it is important to consider that ML architectures also need to handle noise. Thus, the Haralick features analysis is primarily employed to gain a better understanding of the input data and its texture characteristics, rather than being directly used for prediction purposes.

We conducted an analysis of the Haralick texture features specifically for mitochondria across all datasets. Figure 1**b** presents the relationship between entropy value and the measures of contrast, correlation, and angular second moment for each mitochondria instance in the four datasets. Notably, the Perez dataset exhibits considerable deviation from the training OHSU dataset in all four tracked measures, providing an explanation for the observed low precision results on the Perez dataset.

To gain a deeper understanding of the impact of input texture clustering on the models, we examine the internal feature representation of the models. The backbone, a fundamental component in most detection architectures, is responsible for extracting features from input images, which are then used for further detection tasks. We aim to investigate whether this internal representation of a particular object, specifically mitochondria in our case, retains the texture characteristics specific to each dataset. In other words, we explore whether we can discern mitochondria from the Perez dataset distinctly from those in the other datasets based on their internal feature representation.

To examine the internal feature representation of mitochondria, we extract the bounding boxes of mitochondria instances and pass them through a ResNet50 backbone trained as part of the Deformable DETR (DDETR) detection model. The feature maps obtained after the fourth block of ResNet50 are then dimensionally reduced using principal component analysis. Finally, for visualization purposes, the feature maps are further reduced to two dimensions using t-distributed stochastic neighbor embedding (t-SNE).

In the left panel of Figure 3**b**, we can observe that the initial texture clustering of the datasets is preserved by the model. This suggests that the model’s internal feature representation still retains the texture characteristics specific to each dataset.

We conducted four-fold experiments to improve the detectors’ generalizability: 1) Switching to a more complex backbone; 2) Pre-training the backbone on EM-specific data; 3) Direct data augmentation during training; 4) Changing the internal data normalization in the backbone. These experiments aimed to enhance the detectors’ ability to capture intricate features, while mitigating the impact of texture differences.

### Backbone

We investigated the effect of architectural changes and pretraining on the backbone to improve generalizability. Three directions were explored: pre-training, alternative backbones, and deeper networks.

First, we replaced the backbone pre-training on ImageNet with pre-training on the stylized-ImageNet dataset [31], which includes randomly stylized versions of the original images. However, this did not improve performance over the baseline on any of the datasets.

Next, we explored pre-training the backbone on domain-specific data. We experimented with pre-training on unlabeled OHSU data using the SWAV training schema [32] and pre-training on the larger CEM500K dataset [33] using the MoCo-v2 schema [34]. The SWAV pre-training showed some improvements in performance on the OHSU, Lucchi++, and Kasthuri++ datasets but not on Perez. This suggests that the SWAV schema encourages the backbone to learn contrast-invariant features, which benefits the detection task.

Additionally, we considered exchanging the backbone for a deeper version (ResNet101) or more robust alternatives to ResNet50 (such as Res2Net [35] and ResNeXt [36]). However, none of these alternatives outperformed the baseline consistently across all datasets.

Overall, the results presented in Table 2 indicate that unsupervised pretraining on OHSU data using SWAV schema is the most effective approach for enhancing performance on multiple datasets but not on all.

**Table 2.** Studying the effect of architectural change of the backbone of DDETR: AP50.

| Techniques | OHSU DS | Kasthuri++ | Lucchi++ | Perez |
| --- | --- | --- | --- | --- |
| Baseline | 56.6% | 51.4% | 56.7% | 16.3% |
| ResNeXt | 60.8% ↑ | 42.6% ↓ | 57.8% ↑ | 0.0% ↓ |
| ResNet101 | 57.0% ↑ | 35.8% ↓ | 64.8% ↑ | 0.0% ↓ |
| stylized-IN ResNet | 56.1% ↓ | 21.7% ↓ | 53.8% ↓ | 0.0% ↓ |
| Moco CELLEM | 58.7% ↑ | 37.3% ↓ | 27.0% ↓ | 0.0% ↓ |
| SWAV OHSU | 57.4% ↑ | 52.5% ↑ | 60.7% ↑ | 0.0% ↓ |

### Texture Augmentation

To improve generalization, we explore texture augmentation as a method of augmenting the training data. Texture augmentation involves applying photometric transformations that specifically target the texture of the images, such as variations in jitters, contrast, sharpness, noise, blurring, and histogram equalization. These aim to introduce additional plausible texture variability without modifying the object annotations. Examples of texture-augmented images are shown in Figure 2. By incorporating these texture augmentations into the training process, we aim to enhance the models’ ability to generalize to datasets with different texture characteristics.

**Fig. 2.**
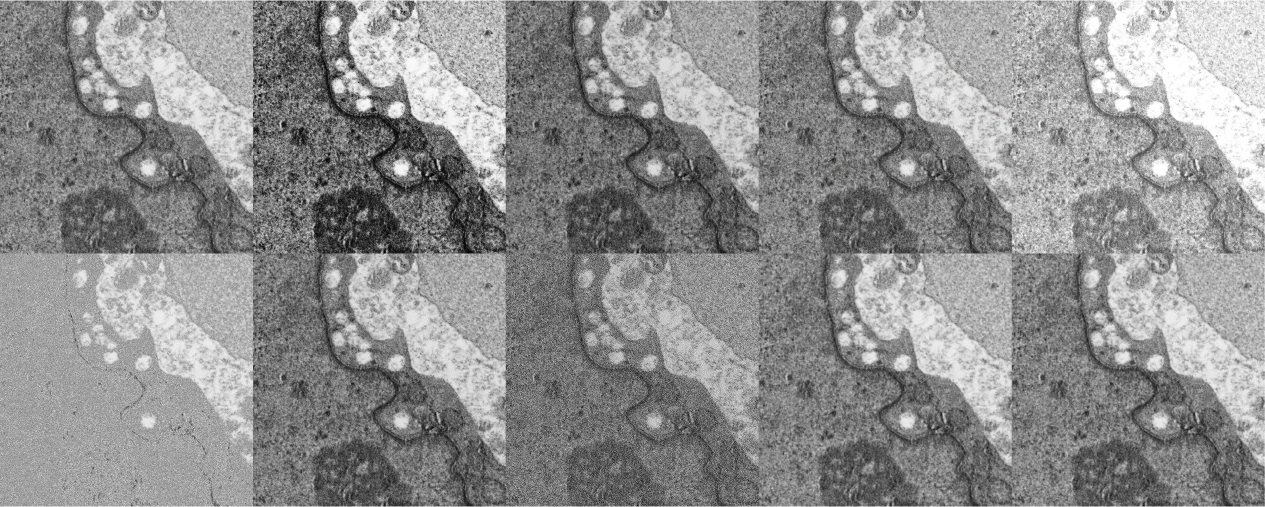
Left to Right: Original, Histogram Equalize, Color Jitter, Gaussian Noise, Random Brightness-Contrast, Random Snow, Hue-Saturation, Sharpen, Blur, Median blur.

The results in Table 3 demonstrate that data augmentation has a significant positive impact on the generalizability of the models. On the unseen data, we observed improvements ranging from 7.5% to 20.5% on average, while on the OHSU test dataset, the average improvement was 3.7%.

**Table 3.** Training using different Intervention techniques - Texture Augmentation, Group Normalization and both. Outcome measured as difference in Average Precision (AP50) from Baseline.

| Intervention | Methods | OHSU DS | Kasthuri++ | Lucchi++ | Perez |
| --- | --- | --- | --- | --- | --- |
| Texture Augmentation | DDETR | +4.1% | +19.4% | +14.9% | 0.0% |
|  | DETR | +1.8% | -0.4% | +4.9% | +27.6% |
|  | Sparse RCNN | +1.7% | -8.8% | +19.4% | +15.4% |
|  | TOOD | +2.8% | +11.9% | +22.3% | +33.6% |
|  | Dynamic RCNN | +7.9% | +15.2% | +20.5% | +11.6% |
|  | YOLOv3 | +3.8% | +7.9% | +18.2% | +30.7% |
|  | Average Diff | 3.7% | 7.5% | 16.7% | 19.8% |
| Group Norm | DDETR | +9.1% | +26.5% | +23.4% | +15.1% |
|  | DETR | +3.3% | +23.1% | +23.5% | +30.3% |
|  | Sparse RCNN | +3.2% | +29.9% | +41.3% | +25.5% |
|  | TOOD | +4.8% | +48.7% | +30.9% | +33.5% |
|  | Dynamic RCNN | +9.1% | +46.5% | +37.8% | +24.5% |
|  | YOLOv3 | +1.4% | +19.0% | +32.2% | +32.9% |
|  | Average Diff | 5.1% | 32.3% | 31.5% | 27.0% |
| Augmentation + Group Norm | DDETR | +8.8% | +4.8% | +20.0% | +6.7% |
|  | DETR | +4.6% | +4.8% | +21.8% | +30.6% |
|  | Sparse RCNN | +3.3% | +11.3% | +38.4% | +25.5% |
|  | TOOD | +7.0% | +59.5% | +27.6% | +15.3% |
|  | Dynamic RCNN | +10.7% | +37.0% | +36.1% | +28.2% |
|  | YOLOv3 | +4.7% | +10.5% | +29.9% | +26.2% |
|  | Average Diff | 6.5% | 21.3% | 29.0% | 22.1% |

It is worth noting that texture augmentation, which was specifically tailored to the OHSU dataset, led to improvements in most cases, except for Sparse RCNN on the Kasthuri++ dataset. This highlights the importance of carefully selecting appropriate augmentation strategies, as unsuitable choices can potentially harm performance, as noted in previous research [13].

To gain a better understanding of how texture augmentation contributes to these improvements, we investigated the internal feature representation of mitochondria using the DDETR architecture.

In Figure 3 **a.**, we observe that the augmented training dataset effectively covers the contrast-entropy space of the Lucchi++ and Kasthuri++ datasets, and even to some extent, the Perez dataset in terms of entropy. This indicates that the texture augmentation strategy successfully introduces additional variability in the training data, allowing the models to learn a wider range of texture patterns.

**Fig. 3.**
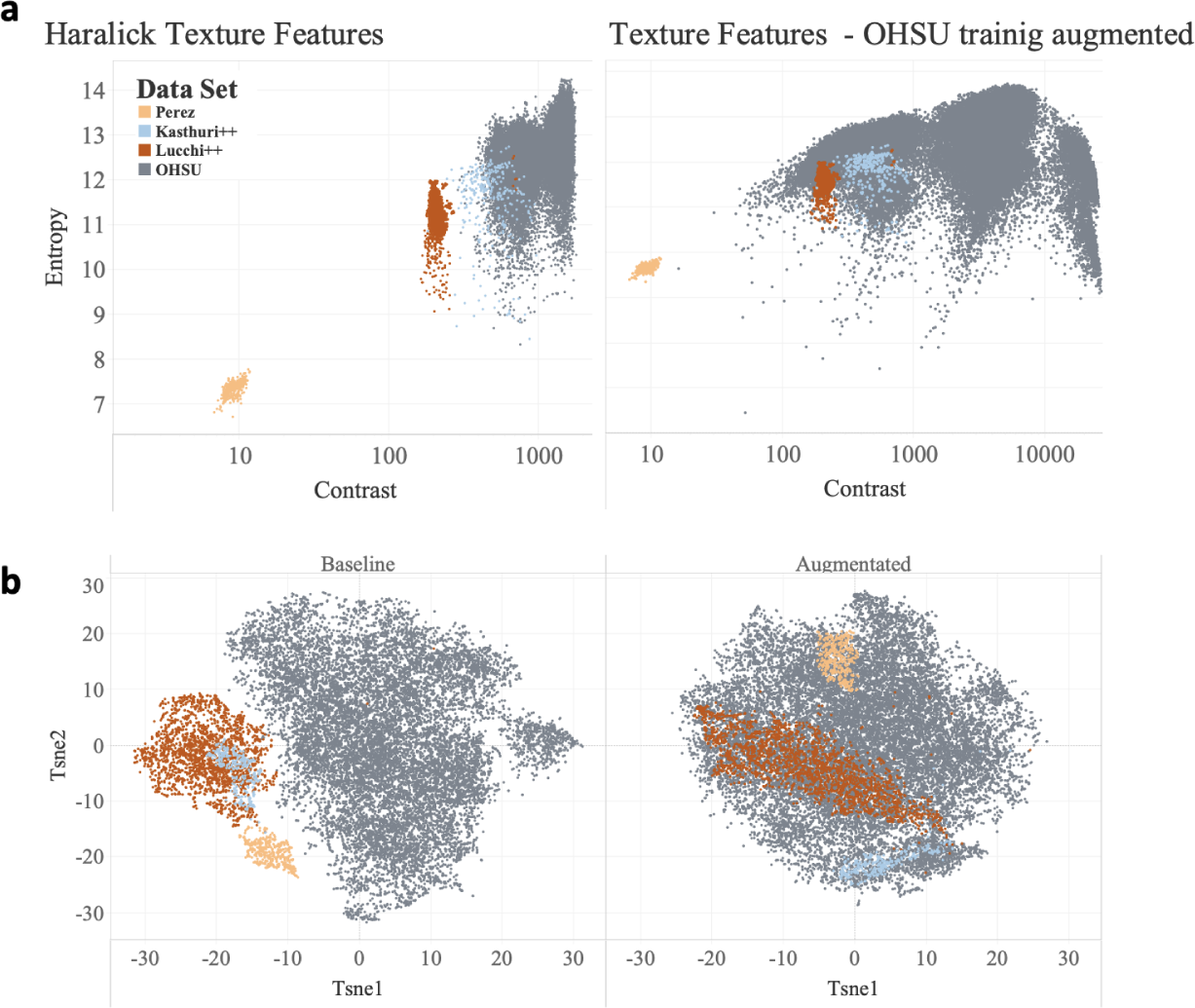
Texture Augmentation: **a.** The Haralick entropy vs. contrast for OHSU training data, Perez, Lucchi++ and Kasthuri++. The left panel depicts the features before augmenting the training dataset and the right panel after the augmentation. **b.** t-SNE visualization of the internal feature representation of the mitochondria crops. The left panel depicts the features of the baseline model; the right panel shows the expansion of the features when the models are trained with texture augmented data.

Figure 3 **b.** demonstrates the clear separation in the internal feature representation of mitochondria between the different datasets. However, we also observe that the augmentation helps in closing the gap between the datasets.

### Group Normalization

The final step towards generalizability is replacing the Batch Normalization layer (BN). Most of the detection algorithms use ResNet as the backbone with a default Batch Normalization as a normalization layer. For BN to work well we need to group the training images into batches of size 32 or more during training (the higher the better) [37]. The traditional BN layer calculates mean and variance within a batch of images, making it sensitive to batch size and unsuitable for small batch sizes commonly used in detection algorithms [19, 38, 39]. Group Normalization (GN) [37], a viable alternative to BN, calculates mean and variance per image, making it suitable for small batch sizes. It strikes a balance between Instance Normalization and Layer Normalization and performs well for small batch sizes and close to BN with large batch sizes. GN normalizes input per layer using mean and standard deviation computed along the spatial dimensions and a group of channels. In our experiments, we set the group size *G* to 32 as suggested in [37]. GN also helps to normalize the contrast information of each image.

We replaced BN with GN in all six architectures and trained them on the OHSU training dataset. The results, as shown in Table 3, demonstrate a remarkable improvement in average precision. The two transformer-based detectors achieve over 80% precision on the Lucchi++ dataset, while all other models also show notable improvements. On the more challenging Kasthuri++ dataset, Deformable DETR achieves a precision of 78%, which is a 26% improvement over the baseline. Overall, the switch to Group Normalization leads to a significant 27-32% improvement on the unseen datasets, highlighting the effectiveness of this normalization approach for enhancing generalizability. The t-SNE analysis reveals that GN is the only technique considered here that effectively blends the texture characteristics in a uniform and indistinguishable manner, contributing to the generalizability of all the methods. This can be observed in the left panel of Figure 4.

**Fig. 4.**
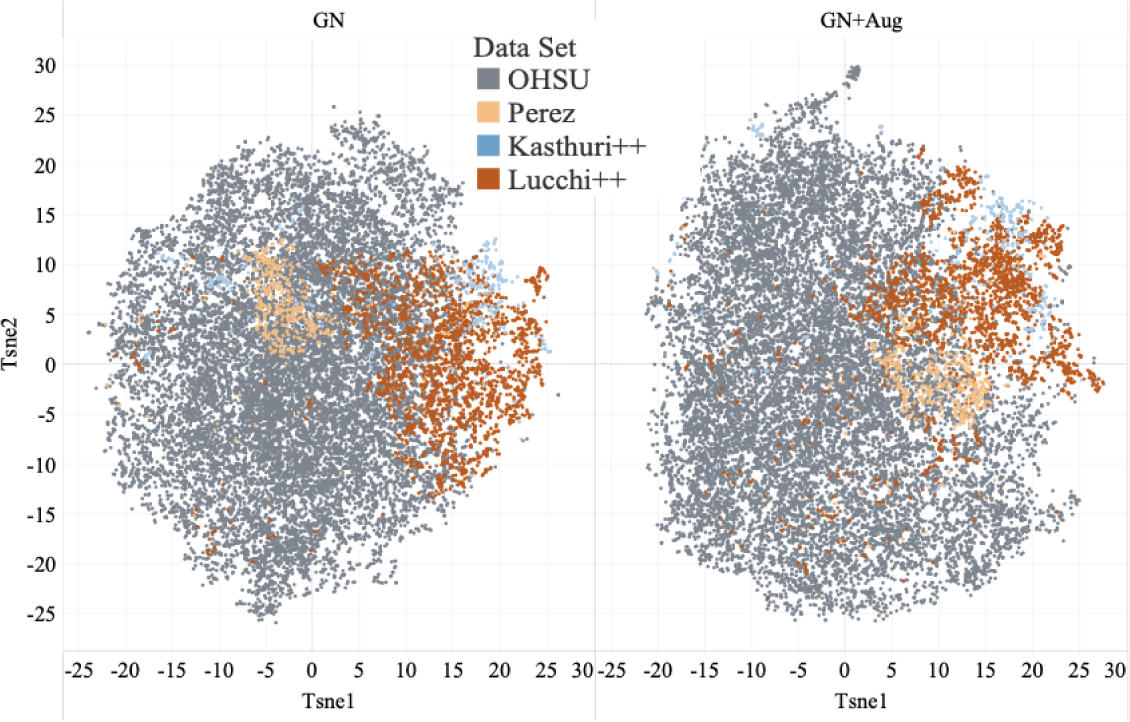
t-SNE feature representation of mitochondria for ResNet50 trained as backbone of DDETR with OHSU training data. **Left panel:** ResNet50 trained with Group Normalization; **Right panel:** ResNet50 trained with Group Normalization and texture augmentation.

Our next step is to combine texture augmentation and group normalization. However, the results in Table 3 demonstrate that applying both strategies only improves performance on the OHSU test dataset. On the other datasets, although there are improvements compared to the baseline, the handpicked texture augmentation actually hinders the performance of GN. This trade-off highlights the balance between task specialization (OHSU data) and expressivity (other datasets). Detection models with stronger structural priors, such as data augmentation, can leverage them to improve precision on specific datasets that benefit from these assumptions. In contrast, models that integrate weaker inductive biases can adapt to diverse domains without relying on restrictive or task-specific conjectures.

## 2 Discussion

Combining data from different sources or research groups, even within the same field, can be challenging due to variations in experimental protocols, sample preparation, equipment, and other factors. This issue extends beyond the biomedical Electron Microscopy field. In this work, we highlight the importance of robustness and generalizability in machine learning techniques and models, emphasizing the need for diverse data. Texture augmentation is presented as a means to diversify the data, but it alone does not ensure inclusiveness.

To address this, we use the Group Normalization layer, which ensures that texturally different data subsets are treated equally by the network. We evaluate the performance of six detection algorithms on four EM datasets, measuring variations in performance. Haralick texture features are used to identify texture differences among the datasets. Despite attempts to address texture clustering through architectural changes, such as increasing backbone size, alternative backbones (ResNext and Res2net), and domain-specific pretraining, the clustering is preserved. Texture augmentation during training encourages the algorithms to learn more diverse features, partially addressing the texture clustering. However, diversity alone does not eliminate differences. The Group Normalization layer is the only technique we found that effectively normalizes the texture representation. It is important to note that this architectural change is independent of the amount or quality of the data. Both texture augmentation and Group Normalization help the models learn features that better generalize to unseen and differently acquired data. We summarize the tested techniques from Tables 2-3 in Table 4, with Group Normalization showing improvements of over 29% and achieving precision rates of over 80% for Lucchi++ and 78% for Kasthuri++. Interestingly, when texture augmentation and Group Normalization are combined, we observe better results only on the OHSU test dataset, with a decrease compared to applying Group Normalization alone.

**Table 4.**
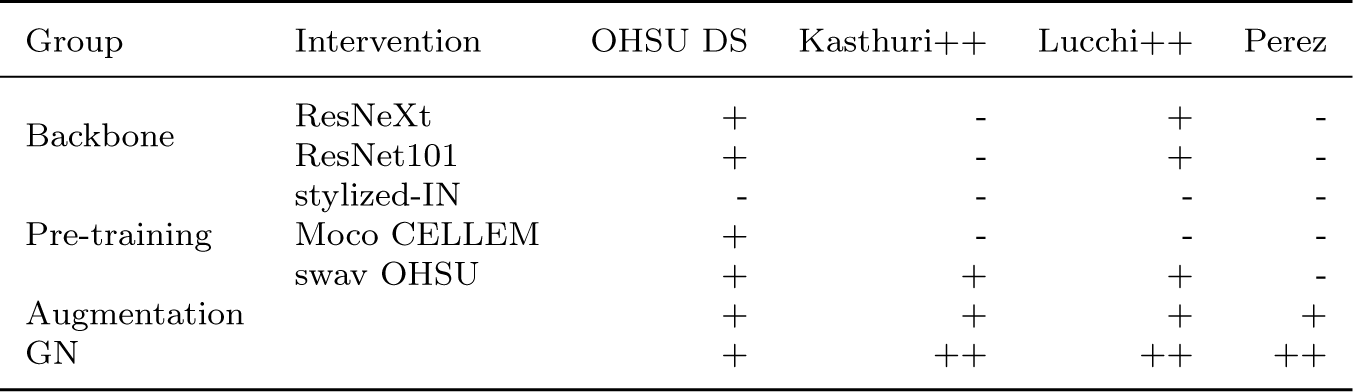
Summary of intervention techniques: ‘+’ positive (1-20%) improvement in performance, ‘-’ zero or negative improvement in performance, ‘++’ over 20% improvement in performance

We plan to further investigate the relationship between data augmentation and Group Normalization, as well as explore new techniques for improving robustness and generalizability in ML models.

## 3 Methods

### OHSU Data collection and Preprocessing

Raw data were collected using an FEI/Thermo Scientific Helios NanoLab DualBeam FIB-SEM microscope equipped with a concentric backscatter detector (CBS) in immersionmode. The Thermo Scientific Maps software package, a commercially-available tool, was utilized for automated collection of large montages of the images.

The OHSU dataset comprises a collection of 527 high-resolution images, each measuring 6*K ×* 4*K*. These images were annotated by 9 annotators, who identified a total of 25,330 endosomes, 30,352 mitochondria, and 903 nucleoli within the images. We used only mitochondria in the current research because that was the only organelle annotated in all of the publicly available datasets (Lucchi++, Kasthuri++ and Perez).

To facilitate training and analysis, the large high-resolution images were sliced into smaller patches measuring 1024 *×* 1024 pixels. The slicing process was performed with a 0.2 overlap ratio, ensuring that adjacent patches overlap by 20% of their width and height. This resulted in a training dataset consisting of 11,520 images, a validation dataset consisting of 1,225 images, and a test dataset consisting of 2,891 images.

During the preprocessing stage, annotations smaller than 20 *×* 20 pixels were removed from the dataset. This step helps to eliminate small or irrelevant annotations that may not provide significant information for the detection models.

### Detection Framework Details

In our research, we utilized the Open MMLab Detection Toolbox repository [19], which is a publicly available collection of detection algorithms and tools. From this repository, we selected six specific architectures to investigate and demonstrate our findings. These architectures were chosen based on their relevance and effectiveness for the task at hand.

| Method | orig config file | Type | anchor<br>Y/N | Back-<br>bone |
| --- | --- | --- | --- | --- |
| Deformable DETR [20] | deformable_detr_twostage_refine_r50_16x2_50e_coco.py | T | N | r50 |
| DETR [21] | detr_r50_8x2_150e_coco.py | T | N | r50 |
| TOOD [23] | tood_r50_fpn_1x_ohsu.py | C | N | r50 |
| Dynamic R-CNN [22] | dynamic_rcnn_r50_fpn_1x_coco.py | C | Y | r50 |
| Sparse R-CNN [24] | sparse_rcnn_r50_fpn_1x_coco.py | C | Y | r50 |
| YOLOv3 [25] | yolov3_d53_mstrain-608_273e_coco.py | C | N | dn53 |

Type T = transformer, C = CNN, r50 = ResNet50, dn53 = DarkNet53

For the datasets used in our research (OHSU, Lucchi, Kasthuri, and Perez), we converted them into COCO format, which is a widely used data format for object detection tasks. The conversion involved modifying the configurations’ data annotation paths and classes to match the specific datasets. Since Lucchi, Kasthuri, and Perez were only used for inference (evaluation), we created additional test dataset configurations for them.

To incorporate texture augmentation into the training process, we added the augmentation techniques to the *train pipeline* section of the configuration files. Specifically, we made use of the Albumentations package integrated into the mmdetection library [40]. Albumentations provides a wide range of image augmentation methods, including photometric transforms, noise addition, blurring, histogram equalization, and others.

Regarding the architectures used, we inherited the published COCO configurations from the Open MMLab Detection Toolbox repository [19]. These configurations served as a starting point for our experiments, and we made necessary modifications such as adjusting the data paths and classes. We initialized the models with the pre-trained weights available from MMLab to benefit from their learned representations.

During training, we mostly followed the original training schedules provided with the COCO configurations, with minor adjustments such as extending the number of training epochs or adapting the learning rate decay steps to suit our specific experiments.

To calculate the Haralick texture features, we utilized the Mahotas image processing library[41].

### Performance Metrics

In a typical detection architecture, the output consists of three predictions: the predicted bounding box coordinates, the confidence score (probability) that the bounding box contains an object, and the class probabilities indicating the likelihood of different object classes.

To evaluate the quality of a prediction, we apply certain criteria and thresholds. The bounding box prediction is used to calculate the intersection over union (IoU) between the predicted box and the ground truth box. IoU measures the overlap between the predicted and ground truth boxes, indicating how well the predicted box aligns with the actual object location.

To determine whether a detection is a true positive (TP), three conditions must be satisfied: 1) The confidence score for the detection is above a specified threshold; 2) The predicted class matches the class of a ground truth object; 3) The IoU between the predicted bounding box and the ground truth box is greater than a specified threshold (e.g., 50%). If any of these conditions are not met, the detection is considered a false positive (FP). If the confidence score for an object is below the threshold, it is considered a true negative (TN), indicating that there is no object present in the predicted bounding box.

Precision is the ratio of TP to the sum of TP and FP, while recall is the ratio of TP to the sum of TP and FN. AP is calculated by averaging precision values across different recall levels and classes. In COCO evaluation, AP is computed by varying the IoU threshold from 0.5 to 0.95 with a step of 0.05. AP50 refers to the average precision at an IoU threshold of 0.5, which is commonly reported.

### Data Augmentation

In Figure 2, we showcase an image from the OHSU dataset before and after applying various texture augmentations individually. The augmentations include Median Blur, Blur, ColorJitter, HueSaturationValue, Sharpen, Histogram Equalization, Random Brightness-Contrast, Random Snow, Color-Jitter group, Gaussian Noise, and Blur group. These augmentations are applied sequentially, with specific probabilities for each augmentation technique. We implemented these augmentations using the Albumentations library [40].

In the case of the YOLOv3 architecture, the original version already incorporates PhotoMetricDistortion as part of its training pipeline, which affects brightness, saturation, hue, and contrast. However, for the “Aug” and “Aug+GN” versions of YOLOv3, we made modifications by removing the Expand augmentation and replacing PhotoMetricDistortion with our texture augmentation technique. This ensures consistency with the texture augmentation applied to the other detection architectures.

## Acknowledgments

The authors acknowledge support from the OHSU Serial Measurements of Molecular Architecture and Response to Therapy (SMMART) clinical trial program, National Institutes of Health (NIH)-National Cancer Institute (NCI) Human Tumor Atlas Network (HTAN) Research Center (U2CCA233280), the Prospect Creek Foundation, OHSU Brenden-Colson Center for Pancreatic Care, a NIH/NCI Cancer Systems Biology Consortium Center (U54CA209988), the OHSU Knight Cancer Institute-Cancer Early Detection Advanced Research (CEDAR) program, and the NIH-Knight Cancer Institute Cancer Center Support Grant (5P30CA69533). Electron microscopy was performed at the OHSU Multiscale Microscopy Core, an OHSU University Shared Resource, with assistance from Erin S. Stempinski and Steven K. Adamou.

## Declarations

This study was approved by the Oregon Health & Science University protocol #16113 Institutional Review Board (IRB). Participant eligibility was determined by the enrolling physician and an informed consent was obtained prior to all study protocol related procedures.

JWG has licensed technologies to Abbott Diagnostics; has ownership positions in Convergent Genomics, Health Technology Innovations, Zorro Bio and PDX Pharmaceuticals; serves as a paid consultant to New Leaf Ventures; has received research support from Thermo Fisher Scientific (formerly FEI), Zeiss, Miltenyi Biotech, Quantitative Imaging, Health Technology Innovations and Micron Technologies; and owns stock in Abbott Diagnostics, AbbVie, Alphabet, Amazon, Amgen, Apple, General Electric, Gilead, Intel, Microsoft, Nvidia, and Zimmer Biomet.

